# Leaky scanning translates a conserved ORF in the 3′UTR of *RPL36A* and regulates the expression of ribosomal protein L36a

**DOI:** 10.64898/2026.05.26.727855

**Authors:** Debraj Manna, Japita Ghosh, Solanki Dhruvi Chetanbhai, Kirtana Vasu, Sandeep M Eswarappa

## Abstract

The 3′UTRs of eukaryotic mRNAs are major hubs of post-transcriptional regulation that govern the spatial and temporal regulation of gene expression. Although 3′UTRs were long considered non-coding, recent studies have revealed translation within downstream open reading frames (dORFs) located in 3′UTRs. In contrast to upstream ORFs (uORFs) in 5′UTRs, the mechanisms governing dORF translation and their functional significance remain poorly understood. Here, we identify and characterize a dORF within the 3′UTR of *RPL36A*, which encodes ribosomal protein L36a. Analysis of ribosome profiling datasets revealed ribosome footprints aligned with the dORF reading frame across multiple human tissues and cell lines. Conservation analysis together with luciferase- and GFP-based reporter assays further supported active translation of the dORF. Using strategically placed RNA hairpin structures and mutational analyses, we show that dORF translation is consistent with ribosomal leaky scanning. Disruption of dORF translation, either by mutating its start codon in exogenous constructs or by CRISPR-mediated deletion of the endogenous dORF-containing region, reduced *RPL36A* mRNA and protein levels and resulted in impaired global translation. We further identified a miR-5701-binding site within the dORF region and provide evidence that dORF translation antagonizes miR-5701-mediated repression of *RPL36A*. Finally, we identify a similar dORF-associated regulatory mechanism in *RPS3,* which encodes ribosomal protein S3. Together, our findings uncover dORF translation as a regulatory mechanism controlling ribosomal protein expression.

## INTRODUCTION

Maintaining the correct stoichiometry of ribosomal proteins (RPs) is essential for efficient ribosome biogenesis and preservation of ribosome integrity. Because ribosome assembly requires near-equimolar production of its components, even modest imbalances can disrupt assembly and trigger cellular stress responses. Excess unassembled RPs can induce cell-cycle arrest, senescence, or apoptosis via MDM2/HDM2-p53 pathway (1-5). Haploinsufficiency of genes encoding specific RPs causes ribosomopathies such as Diamond-Blackfan anemia and 5q-syndrome. These disorders highlight the physiological importance of tight quantitative control of RP expression. Notably, despite the universal requirement for ribosomes, ribosomopathies often exhibit striking tissue-specific phenotypes (e.g., anemia), suggesting that RP gene expression is subject to context-dependent regulatory mechanisms (6,7).

The expression of RPs is regulated at multiple levels to ensure coordinated ribosome biogenesis. A prominent regulatory feature of many RP mRNAs is the 5′-terminal oligopyrimidine (5′TOP) motif, an oligopyrimidine tract located immediately downstream of the mRNA cap that confers sensitivity to the mTOR signaling pathway (8,9). Under favourable conditions, TOP mRNAs are efficiently translated, but stress or mTOR inhibition selectively represses their translation, allowing cells to synchronously dial down ribosome production when nutrients are scarce (10,11). In addition to this translational control mechanism, RP mRNAs are subject to post-transcriptional regulation by microRNAs (miRNAs). Numerous miRNAs have been reported to bind RP transcripts and modulate their expression by influencing mRNA stability or translational efficiency (12,13). In contrast, certain miRNAs can enhance RP mRNA translation; for example, miR-10a binds to sequences downstream of the 5′TOP motif within the 5′ untranslated regions (UTRs) of several RP mRNAs and promotes their translation (14). Together, these regulatory mechanisms allow small non-coding RNAs to contribute to the homeostatic control of RP expression and ribosome biogenesis.

Many mRNAs harbour translatable open reading frames (ORFs) or translons in their UTRs (15-17). Translation in these ORFs is being illuminated by sensitive techniques such as ribosome profiling and mass spectrometry (18-22). Some of them even yield functional micro-peptides (23-26). Upstream translons (i.e., uORFs that are translated) can markedly attenuate translation of the canonical ORF (27,28). Translons in the 3′UTRs (i.e., translatable dORFs) were discovered relatively recently (29-34). Previous studies suggest that their translation also regulates the expression of their corresponding mRNAs (29,35). However, the mechanism by which dORFs are translated is poorly understood, and how dORF translation regulates gene expression remains unknown.

Because dORFs are positioned downstream of canonical coding sequences, non-canonical mechanisms are required for their translation. One such mechanism is ribosomal leaky scanning, in which a fraction of scanning pre-initiation complexes bypasses the upstream start codon and initiates translation at a downstream AUG or a near-cognate start codon (36-40). A second mechanism is termination-reinitiation, in which the post-termination 40S ribosomal subunit remains associated with the mRNA after translation of the upstream ORF and subsequently reacquires the factors required for reinitiation at a downstream start codon when the local sequence context is permissive (41-43). Translation of downstream ORFs can also potentially occur through internal ribosome entry, a cap-independent mechanism in which ribosomes are recruited directly to an internal region of the mRNA. Such initiation is directed by internal ribosome entry sites (IRESs), cis-acting RNA elements that recruit the translation machinery independent of 5′ cap. This mechanism is used by many viral and some cellular mRNAs (44,45).

In this study, we identify and functionally characterize an evolutionarily conserved dORF in *RPL36A*. We show that this dORF is translationally active and that its translation is consistent with a ribosomal leaky scanning mechanism. Our results indicate that the dORF translation antagonizes miR-5701-mediated repression of *RPL36A* mRNA, thereby promoting transcript abundance. Consistent with this regulatory role, mammalian cells lacking the dORF exhibit decreased global protein synthesis. Together, these findings reveal a post-transcriptional regulatory mechanism in which translation of a 3′UTR-embedded dORF modulates canonical gene expression.

## RESULTS

### The 3′UTR of *RPL36A* has an evolutionarily conserved translatable ORF

During our analyses to identify RP genes that could potentially exhibit stop codon readthrough, we observed a seven-codon-long dORF in the 3′UTR of human *RPL36A*. This gene encodes ribosomal protein L36a (or eL42), a component of the 60S ribosomal subunit. Interestingly, the positions of both the start and stop codons of the dORF are highly conserved across mammals including the most primitive prototherians. But the 3′UTR sequence before and after this dORF is not conserved (Fig 1A). The start codon (AUG) initiating the *RPL36A* dORF is embedded within a moderate Kozak context, containing a purine residue at position −3 (adenine) but lacking the guanine at position +4. The canonical start codon, however, has strong Kozak context (adenine at -3 and guanine at +4). The selective conservation of this dORF indicated its functional significance. However, there are a few exceptions where this dORF was not observed in the currently available sequences (e.g, *Carlito syrichta*, the Philippine tarsier).

**Figure 1.**
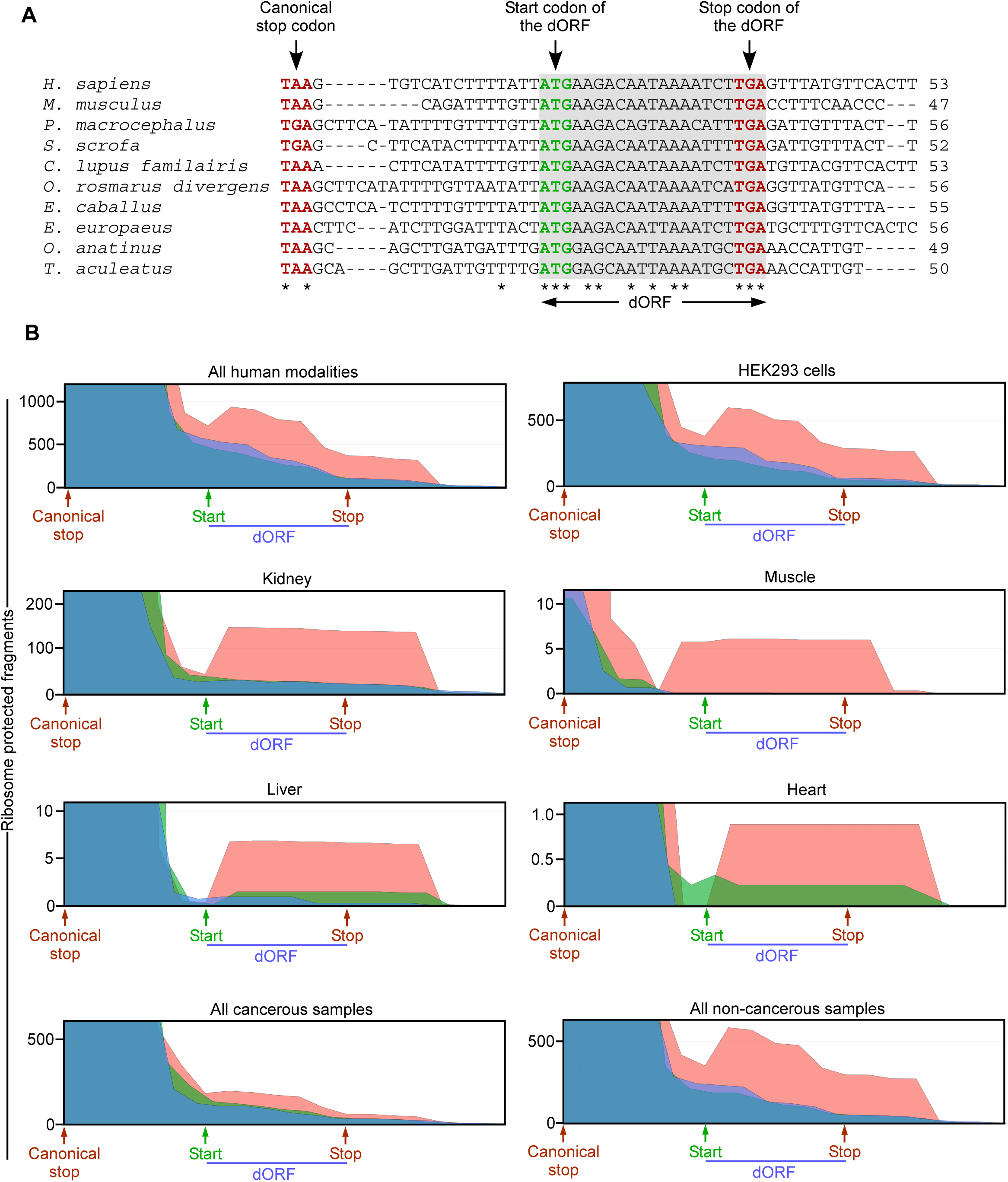
The 3′UTR of *RPL36A* contains an evolutionarily conserved potentially translatable downstream ORF (dORF) (A) Multiple-sequence alignment of the *RPL36A* 3′UTR from representative mammalian species. The stop codon of the canonical ORF and the putative dORF start and stop codons are indicated. Asterisks (*) denote nucleotides that show 100% conservation among the sequences shown. (B) Ribosome-protected fragment (RPF) density plots showing reads uniquely mapped to *RPL36A*. The positions of the canonical stop codon and the dORF are indicated. Plots were generated using the RiboCrypt platform from publicly available ribosome profiling datasets derived from the indicated human cells or tissues. The three colors represent the three different translation frames. The red frame represents the translation frame of the dORF.

Conservation of the start and stop codons suggested that the *RPL36A* dORF is translated. To gather evidence for this, we analysed the ribosome footprints on this dORF using RiboCrypt, a web-based tool for the visualization and analysis of ribosome profiling (Ribo-seq) data. We first analyzed ribosome footprints on the dORF using data from ‘all human modalities’ available in RiboCrypt. We observed a peak in ribosome footprints that started and ended near the start and stop codons, respectively, of the *RPL36A* dORF. Importantly, the peak was observed only in one translational frame, which matched that of the *RPL36A* dORF (Red in Fig 1B). Translation frame of the dORF was inferred based on the translation frame of the canonical ORF. Similar observations were made in Ribo-seq data derived from HEK293 cells, human kidney, liver, and heart tissues. These observations provided evidence for the translation of *RPL36A* dORF both in cell lines and human tissues. Notably, the ribosome footprint peak was absent in cancerous tissue, but present in corresponding non-cancerous control tissue (Fig 1B). This observation suggested regulated translation of *RPL36A* dORF. Furthermore, *RPL36A* dORF was identified as a potential translatable ORF in a previous study. The analysis was done using multiple ribosome sequencing analysis pipelines, including RibORF, Ribo-TISH, and PRICE (30).

### Experimental evidence for the translation of the *RPL36A* dORF

We next performed experiments to test the translatability of the *RPL36A* dORF. The coding sequence (CDS) of *RPL36A*, along with the proximal 33 nucleotides of its 3′UTR, was cloned upstream of the CDS of firefly luciferase (FLuc). The proximal 33 nucleotide stretch of the 3′UTR contains the dORF and the 15 nucleotides between the canonical CDS and the dORF, termed iUTR (internal UTR). The canonical CDS, the dORF, and FLuc were in the same translation frame. The stop codon of the dORF and the start codon of the FLuc were not included, such that FLuc translation is possible only if the dORF is translated (Schematic in Fig 2A). This construct was transfected into HEK293 cells, and the luciferase activity was measured after 24 h. We observed robust luciferase activity, well above that in cells transfected with the construct lacking the proximal 3′UTR. Importantly, mutating the AUG of the dORF to AUC reduced the luciferase activity to background levels. Deletion of the iUTR, which brought the canonical CDS and the dORF next to each other, reduced the luciferase activity but did not abolish it completely. To assess the efficiency of dORF translation, we used another construct in which FLuc was the canonical ORF and therefore translated from the 5′ cap. When transfected into HEK293 cells, this construct showed 3-fold higher luciferase activity than the construct in which FLuc was translated as part of the dORF (Fig 2B). We then subjected the constructs to *in vitro* transcription followed by *in vitro* translation using rabbit reticulocyte lysate. dORF translation was observed in this cell-free assay system, ruling out artifacts such as cryptic promoters and splicing as the reason behind reporter activity (Fig 2C). Collectively, these results support our hypothesis that *RPL36A* dORF is translated and show that the efficiency of its translation in cells is 22.67±4% of the cap-mediated translation of the canonical ORF.

**Figure 2.**
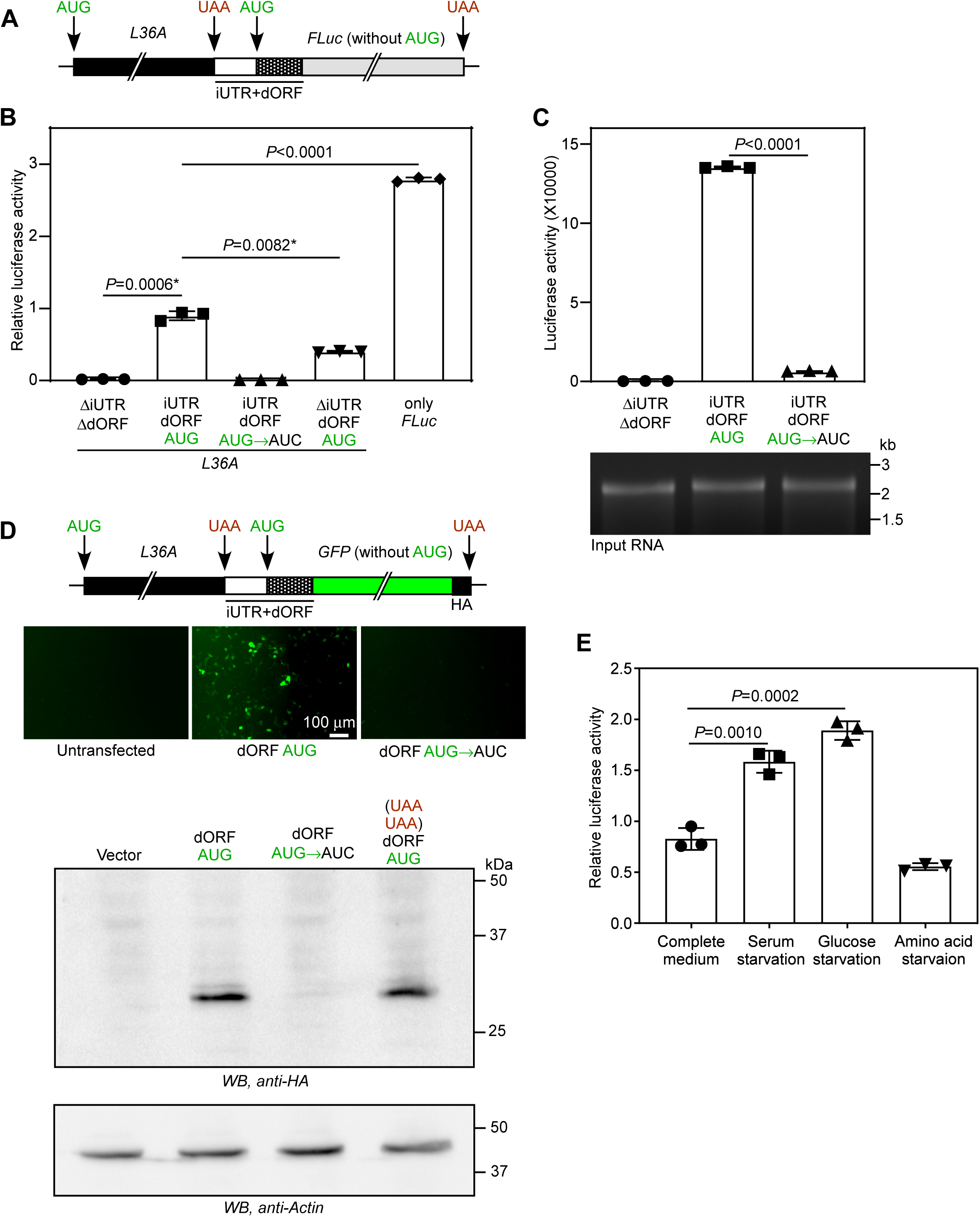
The *RPL36A* dORF is translationally active. (A) Schematic of the firefly luciferase (FLuc) reporter used to assess translation of the *RPL36A* dORF. The canonical *RPL36A* coding sequence together with its 3′UTR, comprising the iUTR (sequence between the canonical ORF and the dORF) and the dORF lacking its native stop codon, was cloned upstream of and in frame with the coding sequence of FLuc lacking its initiation codon, such that luciferase expression depends on translation extending from the dORF. (B) Luminescence-based assay to test the translation of the dORF. Relative luciferase activity in HEK293 cells transfected with the indicated constructs. Firefly luciferase activity was normalized to co-transfected *Renilla* luciferase activity to control for transfection efficiency. (C) Absolute firefly luciferase activity following *in vitro* transcription and translation of the indicated constructs. Translation was performed using rabbit reticulocyte lysate. (D) The schematic shows the GFP-based reporter used to independently investigate translation of the *RPL36A* dORF. Representative fluorescence microscopy images of HEK293 cells transfected with the indicated GFP reporter constructs are shown. Images were acquired using an IXplore IX73 inverted fluorescence microscope with a 10× objective lens. The western blot shows HA-tagged GFP expression in HEK293 cells transfected with the indicated constructs. L36A(UAAUAA)dORF had an additional in-frame stop codon before the dORF. (E) Luminescence-based assay to examine the effect of nutrient stress on dORF translation. Relative luciferase activity was measured in HEK293 cells transfected with the construct shown in (A). Four hours after transfection, the medium was replaced with complete medium or medium lacking serum, glucose, or amino acids. After 20 h of incubation, luciferase activity was quantified. Firefly luciferase activity was normalized to co-transfected *Renilla* luciferase activity to account for differences in transfection efficiency. Bars represent mean ± SD from n = 3. Statistical significance was determined using a two-tailed unpaired Student’s t-test. Data shown are representative of at least three independent experiments. *, Welch’s correction applied.

To further confirm the translation of *RPL36A* dORF, we used green fluorescent protein (GFP) as a reporter. We replaced the CDS of FLuc with that of GFP such that translation of the dORF results in GFP expression and fluorescence, which can be quantified. We observed fluorescence only in cells transfected with a construct having an intact dORF. We also detected GFP protein by western blotting. Mutation of the dORF start codon AUG to AUC abolished the GFP expression and fluorescence (Fig 2D). Together, these observations provide additional evidence for translation of *RPL36A* dORF. We then investigated if the translation of the dORF is a regulated process. Because nutrient stresses activate non-canonical modes of translation (46-49), we tested dORF translation using the construct described in Fig 2A. We observed enhanced translation of the dORF under serum-starvation and glucose-starvation conditions. However, amino acid starvation did not alter dORF translation in this assay (Fig 2E).

### Translation of the *RPL36A* dORF is mediated neither by internal ribosome entry nor by stop-codon readthrough

After demonstrating translation of the *RPL36A* dORF, we next investigated the underlying mechanism. Because this dORF resides within the 3′UTR, downstream of the canonical ORF, its translation must rely on a noncanonical initiation mechanism, such as leaky scanning, termination-reinitiation, or internal ribosome entry.

We tested the *RPL36A* sequence upstream of the dORF for potential IRES activity that can initiate translation of the dORF. For this, we performed the dual-luciferase-based IRES assay. The sequence comprising partial CDS (75 nucleotides) and the iUTR of *RPL36A* was cloned between the CDSs of *Renilla* luciferase (RLuc) and firefly luciferase (FLuc) (schematic in Fig 3A). In this construct, RLuc translation is mediated by the 5′ cap, and FLuc translation is mediated by the IRES activity, if any, of the test sequence. The ratio of FLuc to RLuc activity is used to quantify the IRES activity. When transfected into HEK293 cells, the *RPL36A* sequence (partial CDS and iUTR) did not exhibit IRES activity above the background level shown by a negative control, which had a non-specific sequence as the test sequence. However, the known IRES element from Encephalomyocarditis virus (EMCV) showed IRES activity, which served as the positive control (50)(Fig 3A). Thus, the results of this assay do not support IRES as a mechanism of translation initiation of the dORF in *RPL36A*.

**Figure 3.**
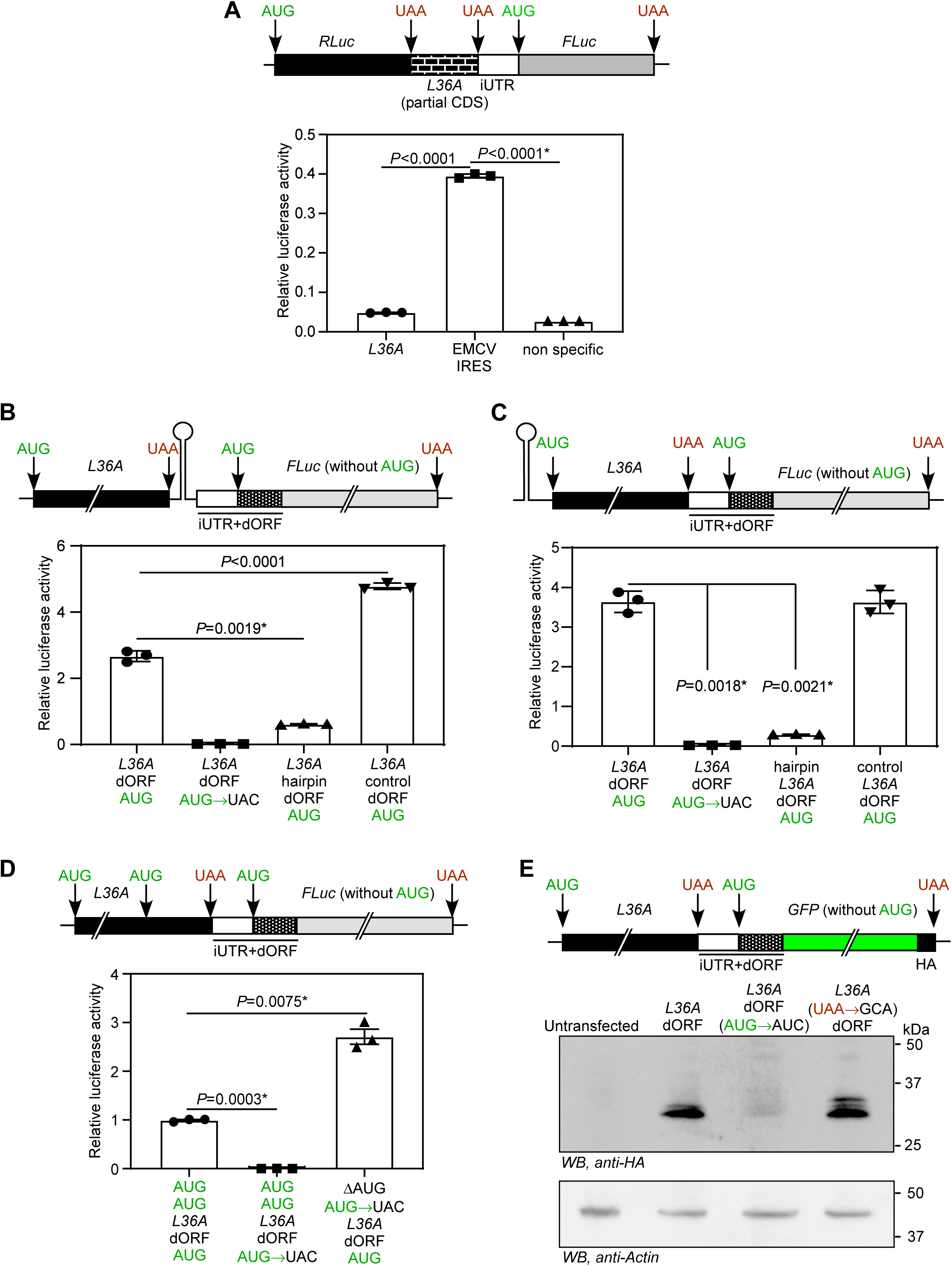
Mechanistic analysis of *RPL36A* dORF translation. **(A)** Dual-luciferase assay used to test internal ribosome entry. HEK293 cells were transfected with the indicated bicistronic reporter constructs expressing *Renilla* luciferase upstream of Firefly luciferase. The graph shows relative luciferase activity (FLuc/RLuc ratio). **(B,C,D)** Luciferase-based assays used to evaluate the contribution of ribosomal scanning-dependent mechanisms for dORF translation. The indicated reporter constructs carrying a stable hairpin-forming sequence positioned either downstream (B) or upstream (C) of the canonical *RPL36A* coding sequence were transfected into HEK293 cells. Two in-frame start codons of the canonical ORF are mutated in the construct used in (D). Firefly luciferase activity was normalized to co-transfected *Renilla* luciferase activity. (E) Western blot analysis used to distinguish between ribosomal leaky scanning and termination-reinitiation mechanisms. Expression of the dORF-derived GFP-HA fusion protein from the indicated constructs is shown. Bars represent mean ± SD from n = 3. Statistical significance was determined using a two-tailed unpaired Student’s t-test. Data shown are representative of at least three independent experiments. *, Welch’s correction applied.

Absence of IRES activity suggested that the dORF is translated from ribosomes that continue from the canonical ORF. One possibility is stop codon readthrough, where ribosomes bypass or miscode the stop codon as a sense codon, resulting in continuation of translation beyond the stop codon (51). However, the western blotting experiments shown in Fig 2C do not support stop codon readthrough as the expected readthrough product of size 43 kDa was not detected. Furthermore, insertion of an extra stop codon after the canonical ORF did not alter the translation of the dORF, which also goes against readthrough as the mechanism (Fig 2D).

### The *RPL36A* dORF is translated through a mechanism consistent with ribosomal leaky scanning

After ruling out readthrough and IRES, we were left with leaky scanning or termination-reinitiation as potential mechanisms. Stable RNA secondary structures, including strong hairpins, are known to inhibit ribosomal scanning by reducing the progression of ribosomal complexes along the mRNA (52,53). Therefore, an RNA hairpin structure can block ribosomal leaky scanning and termination-reinitiation. To test them, we used a 42-nucleotide sequence that forms a stable RNA hairpin structure (54). Using a dual-luciferase construct, we confirmed that this sequence inhibits cap-mediated translation initiation (Fig S1).

We inserted the 42-nucleotide-long hairpin-forming structure after the stop codon of the canonical ORF in the luciferase reporter construct described above (Schematics in Fig 2A and 3B). When transfected into HEK293 cells, this construct resulted in much lower luciferase activity than the control. Another sequence of the same length (42 nucleotides) that does not form any stable structure was cloned after the stop codon of the canonical ORF and used as a control (Fig 3B). Similarly, the hairpin placed upstream of the canonical ORF (i.e., *L36A*) was able to reduce the dORF translation to near-background levels (Fig 3C). These results suggest that the dORF translation is due to either leaky translation or termination-reinitiation mechanism.

To distinguish between leaky scanning and termination-reinitiation as mechanisms of dORF translation, we mutated the two AUG codons present within the canonical *RPL36A* ORF in the luciferase reporter construct. Specifically, the primary start codon was removed, and the internal in-frame AUG was mutated to abolish translation of the canonical ORF. If dORF translation occurred through termination-reinitiation, disruption of canonical ORF translation would be expected to abolish dORF translation. Instead, transfection of this mutant construct into cells resulted in increased dORF translation (Fig 3D). This observation is consistent with a leaky scanning mechanism, as the dORF AUG becomes the first available translation initiation codon in the reporter’s translation frame in the mutant construct.

Next, we used a modified version of the construct shown in Fig. 2D in which the UAA stop codon of the canonical *RPL36A* ORF was mutated to a sense codon, thereby extending translation of the canonical ORF into the downstream region containing the dORF. Under the termination-reinitiation model, elimination of the canonical ORF termination would be expected to abolish the dORF translation because reinitiation requires prior termination of the upstream ORF. In contrast, if the dORF translation occurs via leaky scanning, the dORF product should still be detected, since initiation at the downstream start codon does not depend on upstream translation termination. Upon transfection into HEK293 cells, we detected the dORF translation product irrespective of whether the canonical UAA stop codon was present or mutated (Fig. 3E). These observations are therefore consistent with ribosomal leaky scanning as the mechanism underlying the dORF translation.

### dORF translation positively regulates *RPL36A* mRNA levels

The evolutionary conservation of the dORF, despite limited overall sequence conservation of the surrounding 3′UTR (Fig 1A), suggests that this element is under selective constraint and is therefore likely to have functional significance. Since both uORFs and dORFs have been reported to modulate expression of their corresponding canonical ORFs (28,29,41,55), we examined whether translation of the *RPL36A* dORF influences expression of the canonical ORF (i.e., *RPL36A* itself). To test this, we expressed HA-tagged RPL36A together with its native 3′UTR, including the iUTR, the dORF, and the remaining downstream 3′UTR sequence, in HEK293 cells (Fig 4A). In another construct, the dORF initiation codon (AUG) was mutated to UAC to prevent dORF translation without altering the overall 3′UTR architecture. Cells transfected with the dORF-deficient construct showed reduced exogenous *RPL36A* expression at both protein and mRNA levels compared with cells expressing the construct containing an intact translatable dORF (Fig. 4B,C). These observations suggest that translation of the dORF positively influences *RPL36A* expression, likely through an effect on mRNA abundance or stability.

**Figure 4.**
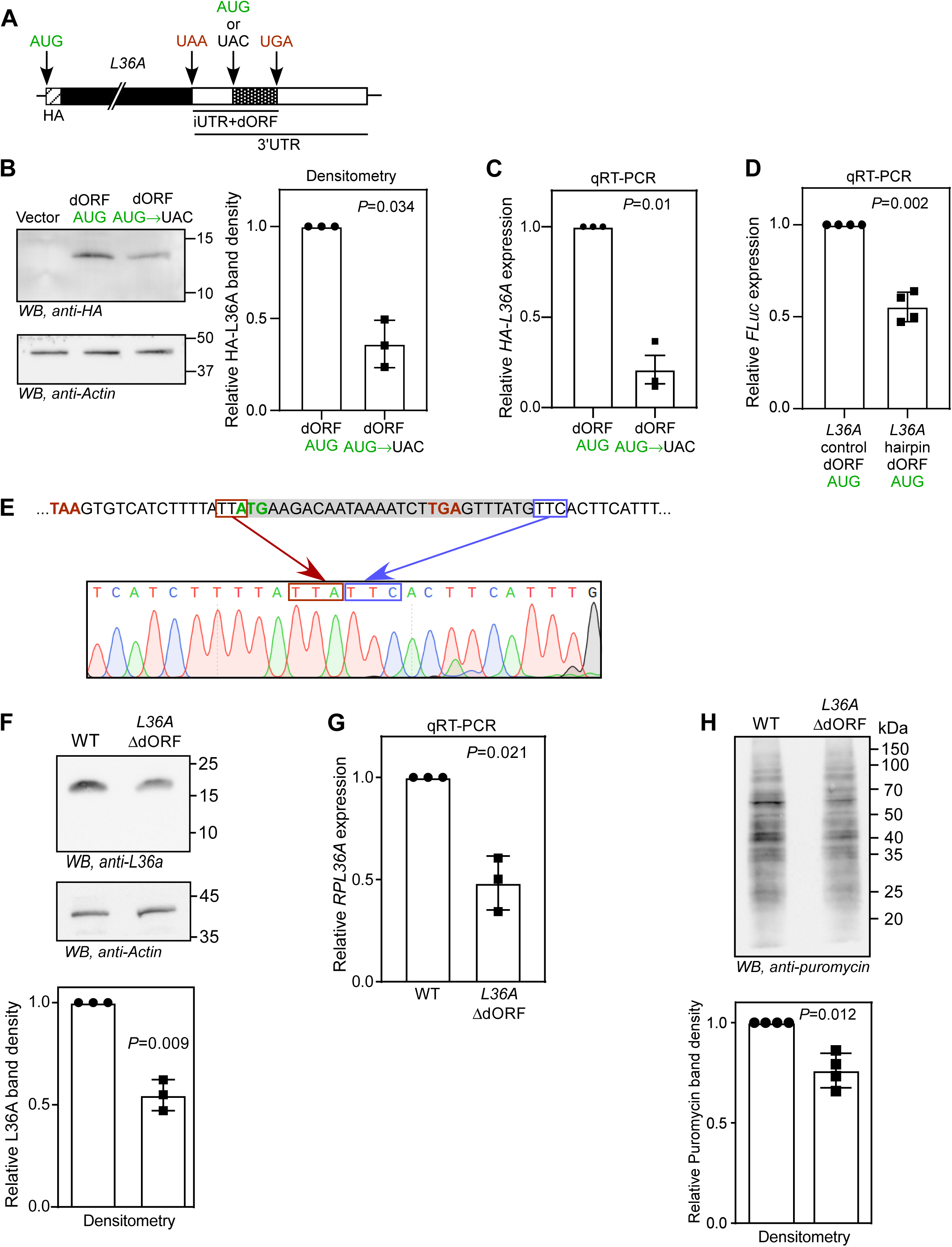
Translation of the *RPL36A* dORF positively regulates *RPL36A* mRNA abundance. (A) Schematic of the reporter construct used to investigate the effect of the dORF translation on *RPL36A* expression. The HA-tagged *RPL36A* coding sequence was cloned together with its native 3′UTR, including the iUTR and the dORF. **(B,C)**Expression of HA-tagged RPL36A in HEK293 cells transfected with the indicated constructs, measured at the protein level by western blotting (B) and at the mRNA level by qRT-PCR (C). In (B), the graph shows densitometric quantification of HA-RPL36A normalized to β-actin. In (C), mRNA levels were normalized to *ACTB*. (D) qRT-PCR analysis of mRNA derived from the luciferase reporter constructs shown in Fig. 3B, comparing constructs with or without the inserted stable hairpin. Transcript levels were normalized to *ACTB*. (E) Sequence of the proximal *RPL36A* 3′UTR, including the iUTR and dORF, indicating the segment deleted by CRISPR-Cas9 in the ΔdORF HeLa clone. Representative sequencing chromatograms confirming the deletion are shown. (F, G) Expression of endogenous *RPL36A* in wild-type and ΔdORF HeLa cells measured at the protein level by western blotting (F) and at the mRNA level by qRT-PCR (G). In (F), the graph shows densitometric quantification of *RPL36A* normalized to β-actin. In (G), mRNA levels were normalized to *ACTB*. (H) Results of the ribopuromycylation assay performed using equal numbers of wild-type and ΔdORF HeLa cells. Cells were treated with 91 µM puromycin for 10 min prior to analysis to assess global translational activity. In all graphs, bars represent mean ± SD from n = 3. Data shown are representative of at least three independent experiments. Statistical significance was determined using a two-tailed paired Student’s t-test.

To further substantiate this observation, we used the hairpin-containing construct described in Fig. 3B, in which insertion of a stable RNA hairpin downstream of the canonical ORF markedly reduces dORF translation (Fig. 3B). When transfected into cells, the hairpin-containing construct exhibited lower mRNA abundance than the control construct carrying a sequence of same length that does not form a stable secondary structure (Fig. 4D). This result is consistent with the idea that suppression of dORF translation is associated with reduced transcript levels.

To determine whether this phenomenon also operates in the endogenous context, we deleted the dORF-encoding region from the endogenous *RPL36A* locus in HeLa cells using the CRISPR-Cas9 system (Fig. 4E). Cells carrying this deletion exhibited reduced endogenous *RPL36A* expression at both protein and mRNA levels compared to parental wild-type cells (Fig. 4F,G). Given that L36a is a constituent of the 60S large ribosomal subunit, we next tested whether reduced L36a abundance affects global protein synthesis. We performed a ribopuromycylation assay. Here, puromycin, an aminoacyl-tRNA analog, is incorporated into elongating nascent polypeptides, enabling quantification of ongoing translation by western blotting with an anti-puromycin antibody. When performed using an equal number of cells, *RPL36A* ΔdORF cells showed reduced puromycin incorporation relative to parental wild-type cells, consistent with decreased global protein synthesis. These findings suggest that loss of the dORF diminishes translational capacity, likely by reducing the availability of L36a. Together, these findings indicate that the dORF contributes to maintaining *RPL36A* mRNA abundance and thereby positively influences global translation.

We then investigated whether the peptide derived from the dORF, MKTIKS, has any detectable function. For this, we used the synthetic peptide MKTIKS and its scrambled version, KTKIMS. HEK293 and HeLa cells were incubated with these peptides with a peptide transfection reagent (for intracellular effects, if any) or without it (for extracellular effects, if any). There was no detectable difference in the morphology or proliferation of cells (Fig S2).

### dORF translation protects *RPL36A* mRNA from miRNA-5701

We next investigated how the translation of the dORF regulates *RPL36A* mRNA levels. We observed the polyadenylation signal, AAUAAA, conserved within the dORF (Fig 1A). The translation of the dORF can potentially mask this signal and inhibit polyadenylation, or the polyadenylation complex can prevent the translation of the dORF. However, canonical polyadenylation occurs in the nucleus, and translation is a cytoplasmic process. Therefore, it is unlikely that translation of the dORF affects the polyadenylation of the mRNA or vice versa. In agreement with this, our polyadenylation assays did not reveal any differences in the polyadenylation of exogenous wild-type vs dORF mutant (AUG to UAC) *RPL36A* mRNAs (Fig S3).

We then analysed the *RPL36A* dORF sequence for potential miRNA targeting sites using miRDB web tool (target score > 75)(56). The analysis revealed that the dORF is a potential target for hsa-miR-5701 (Fig 5A). The target sequence, GACAAUAA, is also conserved in most mammals, suggesting its functional significance (Fig 1A). We confirmed the expression of miR-5701 in HEK293 cells using the miTED web tool and experimentally by RT-PCR (Fig 5B and C)(57). To examine whether miR-5701 regulates *RPL36A* expression, we modulated intracellular levels of this miRNA in HEK293 cells using a miRNA mimic and an inhibitor. Cells transfected with the miR-5701 mimic showed reduced *RPL36A* expression, whereas inhibition of endogenous miR-5701 led to increased *RPL36A* expression. *FGFR2*, a known target of miR-5701, was included as a positive control and showed the expected reciprocal response (Fig. 5D and E)(58). These findings indicate that *RPL36A* expression is negatively regulated by miR-5701.

**Figure 5.**
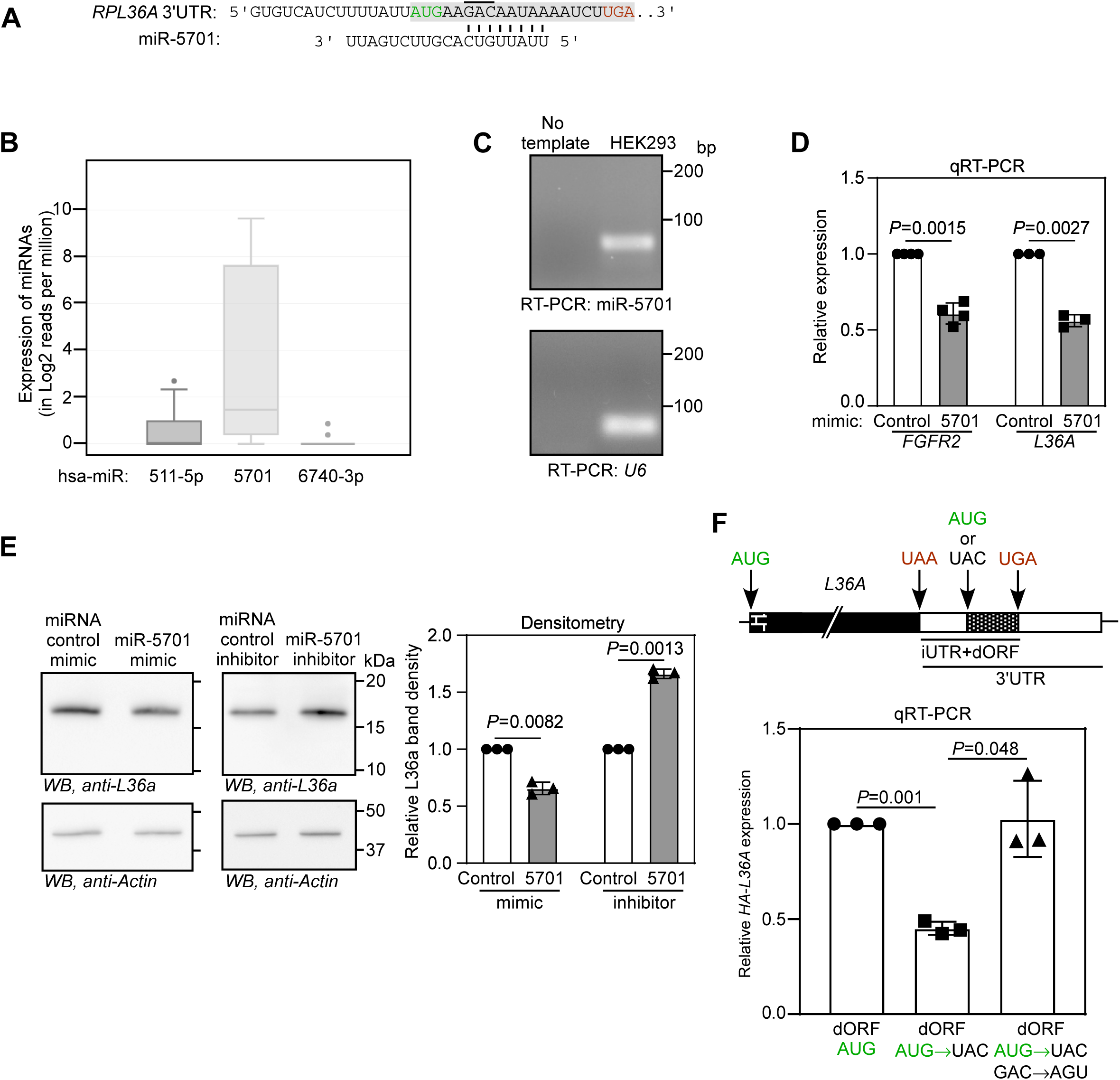
Translation of the *RPL36A* dORF counteracts miR-5701-dependent repression. (A) Sequence alignment of the proximal *RPL36A* 3′UTR, including the iUTR and dORF, with miR-5701. The predicted miR-5701 seed-matching region is indicated. The GAC trinucleotide mutated in the reporter construct used in (F) is highlighted with a horizontal line. (B) Expression profiles of the indicated microRNAs in HEK293 cells, obtained from the miTED online resource. Box-and-whisker plots indicate the minimum, first quartile, median, third quartile, and maximum values. (C) RT-PCR analysis confirming expression of miR-5701 in HEK293 cells. Data shown are representative of three independent experiments. (D) qRT-PCR analysis of endogenous *FGFR2* and *RPL36A* mRNA levels in HEK293 cells transfected with control oligonucleotide or miR-5701 mimic. Transcript levels were normalized to *ACTB*. (E) Western blot analysis of endogenous *RPL36A* expression in HEK293 cells transfected with control oligonucleotide, miR-5701 mimic, or miR-5701 inhibitor. The graph shows densitometric quantification of L36a protein normalized to β-actin. (F) qRT-PCR analysis of exogenous *RPL36A* transcripts in HEK293 cells transfected with the indicated constructs containing the *RPL36A* wild-type sequence or a dORF start-codon mutation, or a mutation disrupting the predicted miR-5701 binding site and dORF start-codon mutation. Transcript levels were normalized to *ACTB*. Bars in all graphs represent mean ± SD, n = 3. Statistical significance was determined using a two-tailed paired Student’s t-test. Data shown are representative of at least three independent experiments.

To investigate whether miR-5701 contributes to the effect of the dORF translation on *RPL36A* mRNA abundance, we used the HA-tagged RPL36A reporter construct described in Fig. 4A. As observed previously, mutation of the dORF initiation codon (AUG), which prevents dORF translation, reduced HA-*RPL36A* mRNA levels. Notably, this reduction was abolished when the predicted miR-5701 binding site was disrupted by point mutation (GAC to AGU; Fig. 5E). The mutations were introduced such that no new miRNA binding sites or protein (known) binding sites are created. These findings indicate that translation of the dORF antagonizes miR-5701-dependent repression, thereby protecting *RPL36A* mRNA from miRNA-mediated downregulation.

### *RPS3* also exhibits regulation by a translatable dORF

We manually screened other mRNAs encoding RPs for conserved dORFs with ribosomal footprints as described above. We identified a conserved dORF within the 3′UTR of *RPS3* (also known as uS3), which encodes the small-subunit ribosomal protein S3. Analysis using the RiboCrypt tool revealed ribosome footprints mapping to the dORF in the same translational frame, consistent with active translation of the dORF. Such footprints were detected across multiple tissues, including embryonic tissue, pancreas, testes, and breast tissue (Fig. 6A,B). We next performed a luminescence-based reporter assay in HEK293 cells, similar to that used for *RPL36A*, to evaluate the translatability of the *RPS3* dORF. Consistent with active translation of the dORF, robust luminescence activity was observed. Importantly, this signal was reduced to near-background levels upon mutation of the dORF start codon to UAC, indicating that the luminescence activity was dependent on translation initiation from the dORF start codon (Fig. 6C). To investigate the effect of the dORF on *RPS3* expression, we cloned either the wild-type or mutant dORF (AUG-to-UAC substitution) downstream of HA-tagged S3 and examined S3 expression in transfected cells. Similar to our observations with *RPL36A*, disruption of dORF translation by mutation of the start codon resulted in reduced expression of HA-tagged S3 compared to the wild-type dORF construct. This reduction was evident at both the protein and mRNA levels (Fig 6D). These results show that S3 expression is also regulated by translation of a dORF.

**Figure 6.**
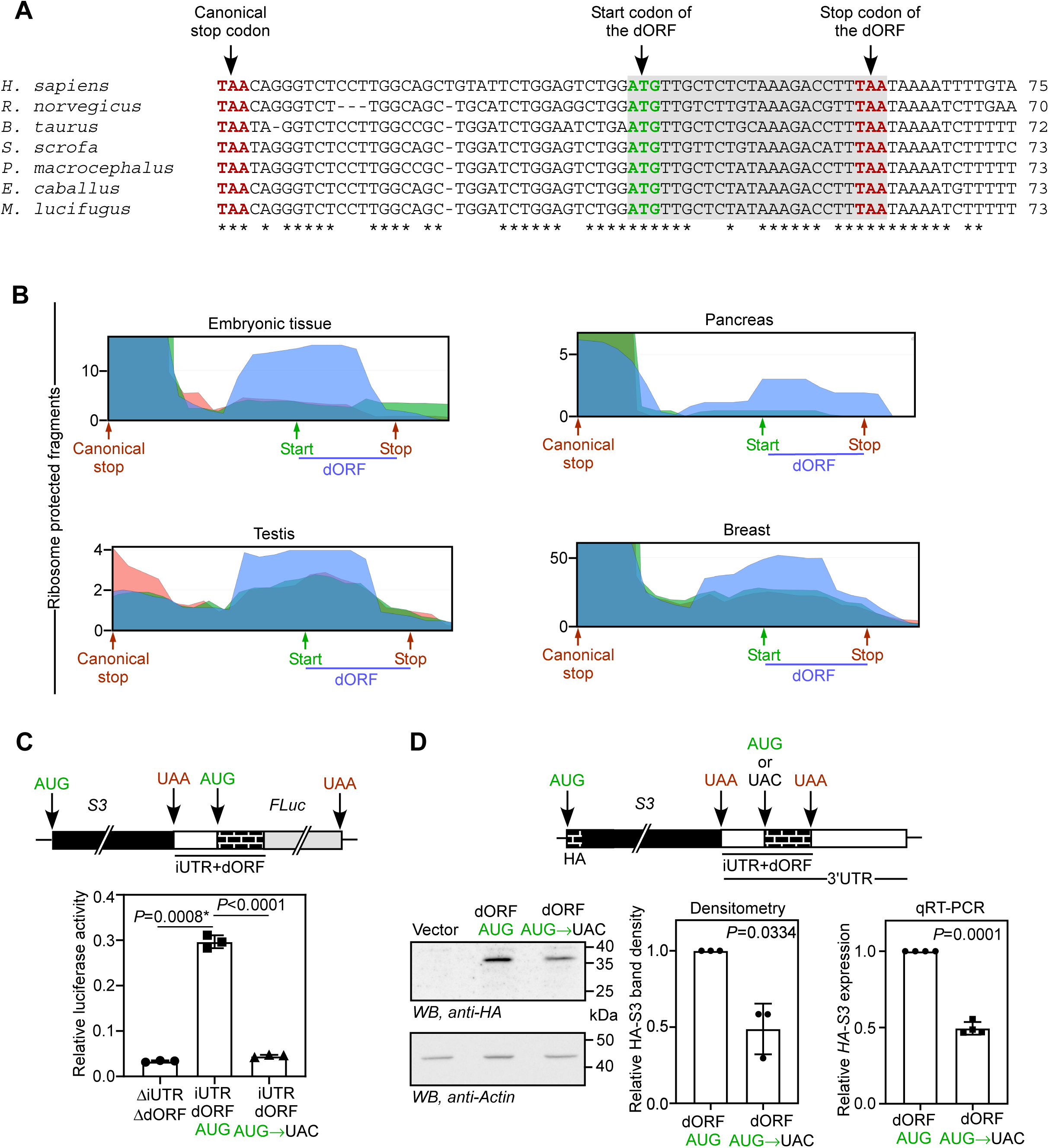
The 3′UTR of *RPS3* contains an evolutionarily conserved translatable dORF. (A) Multiple-sequence alignment of the *RPS3* 3′UTR from representative mammalian species. The canonical stop codon of the canonical ORF and the putative dORF start and stop codons are indicated. Asterisks (*) denote nucleotides that show 100% conservation among the sequences shown. (B) Ribosome-protected fragment (RPF) density plots showing reads uniquely mapped to *RPS3*. The positions of the canonical stop codon and the dORF are indicated. Plots were generated using the RiboCrypt platform from publicly available ribosome profiling datasets derived from the indicated human tissues. Blue represents the translation frame of the dORF. (C) The schematic shows the firefly luciferase (FLuc) reporter construct used to assess translation of the *RPS3* dORF. The canonical *RPS3* coding sequence together with its 3′UTR, comprising the iUTR (sequence between the canonical ORF and the dORF) and the dORF lacking its native stop codon, was cloned upstream of and in frame with the coding sequence of FLuc lacking its initiation codon, such that luciferase expression depends on translation extending from the dORF. The graph shows the results of luminescence-based assay to test the translation of the dORF. Relative firefly luciferase activity in HEK293 cells transfected with the indicated constructs. Firefly luciferase activity was normalized to co-transfected *Renilla* luciferase activity to control for transfection efficiency. (D) The schematic shows the reporter construct used to investigate the effect of dORF translation on *RPS3* expression. The HA-tagged *RPS3* coding sequence was cloned together with its native 3′UTR, including the iUTR and the dORF. Western blot shows the expression of HA-tagged RPS3 in HEK293 cells transfected with the indicated constructs. The densitometry graph shows quantification of HA-RPL36A normalized to β-actin. The qRT-PCR graph shows mRNA levels normalized to *ACTB*. Bars in all graphs represent mean ± SD, n = 3. Statistical significance was determined using a two-tailed unpaired (C) or paired (D) Student’s t-test. Data shown are representative of three independent experiments.

## DISCUSSION

3′UTRs play central roles in the post-transcriptional regulation of gene expression. They contain binding sites for miRNAs and RNA-binding proteins that influence mRNA stability, localization, and translational efficiency, thereby contributing to the spatial and temporal control of protein synthesis (59). Notably, the average length of 3′UTRs increases with organismal complexity, suggesting an evolutionary expansion of regulatory potential in higher eukaryotes (60). In many mammalian mRNAs (e.g., *VEGFA*), the 3′UTR is substantially longer than the coding sequence, suggesting that these regions may mediate multiple layers of post-transcriptional control that remain incompletely understood. For many years, 3′UTRs were considered exclusively non-coding regions. However, advances in high-resolution techniques such as ribosome profiling and mass spectrometry have revealed that translation can occur within 3′UTRs of specific mRNAs, either through stop codon readthrough or via downstream open reading frames (dORFs) (29,51). In this study, we identify a conserved dORF within the 3′UTR of *RPL36A* and demonstrate that its translation regulates the expression of the ribosomal protein L36a.

Multiple lines of evidence support the translation of the *RPL36A* dORF. Analysis of publicly available ribosome profiling datasets using the RiboCrypt platform revealed distinct ribosome footprints aligned with the reading frame of the dORF. Importantly, these footprints were detected across multiple human tissues, indicating *in vivo* translation of the dORF. Consistent with these observations, a previous study independently predicted the *RPL36A* dORF as a translatable ORF using several ribosome profiling-based analytical pipelines (30). In addition, we experimentally validated the dORF translation using luciferase- and GFP-based reporter assays. Interestingly, the dORF translation was enhanced under serum starvation and glucose deprivation, suggesting that its translation may be dynamically regulated in response to nutrient stress.

Because the *RPL36A* dORF is located downstream of the canonical ORF, its translation must occur through a non-canonical mechanism. Potential mechanisms include leaky scanning, termination-reinitiation, and internal ribosome entry. Whereas leaky scanning and termination-reinitiation are dependent on 5′ cap-mediated scanning, internal ribosome entry occurs independently of the 5′ cap. Our dual-luciferase IRES assays did not support the presence of internal ribosome entry activity within the tested *RPL36A* sequence. Insertion of a stable hairpin structure upstream of the dORF abolished its translation, indicating that the dORF translation requires 5′ cap-dependent ribosomal scanning. These observations narrowed the possible mechanisms to leaky scanning or termination-reinitiation. Mutational analysis of the canonical ORF start codon or stop codon supports leaky scanning as the primary mechanism driving translation of the *RPL36A* dORF. Importantly, the dORF translation was also observed in rabbit reticulocyte lysate-based *in vitro* translation assays, ruling out contributions from artifacts such as cryptic promoter activity or alternative splicing.

Because translation is an energetically demanding process, the translation of the *RPL36A* dORF suggests potential functional relevance. The dORF encodes a short six-amino-acid peptide. Interestingly, although the presence of the dORF within the 3′UTR is evolutionarily conserved, the encoded amino acid sequence itself shows limited conservation across species. This observation argues against a conserved peptide-dependent function and raises the possibility that the regulatory role may instead reside in the process of translation itself. Consistent with this interpretation, we did not detect measurable cellular effects from intracellular or extracellular treatment with the synthetic dORF-derived peptide. However, our assays examined only a limited set of cellular conditions and readouts, and it remains possible that the peptide exerts context-dependent functions in specific cell types, physiological states, or stress conditions that were not evaluated in the present study.

Because the canonical polyadenylation site is present within the dORF, we initially considered the possibility that dORF translation might influence polyadenylation of the *RPL36A* transcript. However, polyadenylation occurs co-transcriptionally in the nucleus, whereas dORF translation takes place in the cytoplasm after mRNA export, making a direct mechanistic coupling between these processes unlikely. Consistent with this interpretation, we did not detect any appreciable differences in the polyadenylation profiles of transcripts containing or lacking the dORF.

Translation is increasingly recognized as an important determinant of mRNA stability and turnover (61,62). We therefore investigated whether translation of the *RPL36A* dORF influences mRNA levels. Disruption of the dORF translation, either by mutating its start codon in exogenous reporter constructs or by CRISPR-mediated deletion of the dORF-containing region in the endogenous *RPL36A* locus, resulted in reduced *RPL36A* mRNA and protein levels. These findings suggest that the dORF translation positively regulates *RPL36A* expression, potentially by enhancing mRNA stability. Consistent with the reduced levels of the ribosomal protein L36a, cells carrying the dORF deletion also exhibited decreased global translational output, as assessed by ribopuromycylation assays.

RNA-binding proteins and miRNAs are major regulators of mRNA stability and expression in cells. Interestingly, we identified a putative miR-5701-binding site within the *RPL36A* dORF region. Our experimental analyses confirmed that miR-5701 modulates *RPL36A* expression under the conditions tested, raising the possibility that the dORF translation may interfere with miRNA-mediated repression. Consistent with this model, mutational analyses targeting either the dORF start codon or the predicted miR-5701-binding site supported a functional interplay between dORF translation and miRNA-dependent regulation.

Notably, unlike the *RPL36A* dORF, which is evolutionarily conserved across mammals, miR-5701 is not broadly conserved and is primarily restricted to primates. This evolutionary distinction suggests that the conserved function of the dORF is unlikely to be solely dependent on miR-5701-mediated regulation. In fact, two other miRNAs (miR-511-5p and miR-6740-3p) binding sites are present within the dORF, although with a lower score (<75). Therefore, dORF translation may have additional regulatory roles that remain to be elucidated.

Genes encoding ribosomal proteins are often coordinately regulated through shared regulatory mechanisms, such as the 5′TOP motif and miR-10a (9,14). Such synchronous mechanisms ensure proper stoichiometric assembly of the ribosome. Our findings raise the possibility that translation of dORFs may represent an additional layer of post-transcriptional regulation for RP genes. Consistent with this idea, we observed evidence for a similar dORF-associated regulatory mechanism in *RPS3*. miR-Let7c-3p binding site (Target score 87) is close to its dORF (8 nucleotides downstream of the stop codon), suggesting functional interaction between translation machinery and the RNA-induced silencing complex.

dORF-mediated regulation could also contribute to ribosome heterogeneity by differentially modulating the expression of ribosomal protein paralogs, such as *RPL36A* and *RPL36AL* (63). Such selective regulation may influence ribosome composition and translational specialization under specific physiological conditions. Although definitive extra-ribosomal functions for L36a have not been established, increased *RPL36A* expression has previously been linked to enhanced proliferation and cancer progression (64,65), supporting the biological relevance of its precise regulation.

## STAR Methods

### Nucleotide sequence alignment

This was performed using Clustal Omega. The 3′UTR sequences of *RPL36A* analyzed in this study were obtained from the following species and accession numbers: *Homo sapiens* (NM_021029), *Mus musculus* (NM_019865), *Physeter macrocephalus* (XM_007119008), *Sus scrofa* (XM_021079666), *Canis lupus familiaris* (XM_022416075), *Odobenus rosmarus divergens* (XM_004417743), *Equus caballus* (XM_001492629), *Erinaceus europaeus* (XM_060183411), *Ornithorhynchus anatinus* (XM_029067023), and *Tachyglossus aculeatus* (XM_038748081). The 3′UTR sequences of *RPS3* were obtained from *Homo sapiens* (NM_001005), *Rattus norvegicus* (NM_001009239), *Bos taurus* (NM_001034047), *Sus scrofa* (NM_001044601), *Physeter macrocephalus* (XM_007101443), *Equus caballus* (XM_001494844), and *Myotis lucifugus* (XM_006102877).

### Analysis of ribosome footprints

Ribosome footprint (RFP) profiles on human *RPL36A* (transcript ID: ENST00000553110) and *RPS3* (transcript ID: ENST00000531188) mRNAs were analyzed using RiboCrypt. RFP datasets were selected as the library type for all analyses. For *RPL36A*, the following experiments were analyzed: all_merged-Homo_sapiens_modalities, all_merged-HEK293, all_tissues_Homo_sapiens (with RFP_kidney, RFP_muscle, RFP_liver, and RFP_heart libraries), and all_cancers_Homo_sapiens (with RFP_FALSE and RFP_TRUE libraries representing non-cancerous and cancerous samples, respectively). For *RPS3*, the all_tissues_Homo_sapiens experiment was analyzed using the RFP_embryonic, RFP_pancreas, RFP_testis, and RFP_breast libraries. Analysis parameters were maintained at their default settings. The display mode for reading frames was set to “area” for all analyses.

### Construction of plasmids

The coding sequence (CDS) of *RPL36A* was cloned between the HindIII and BamHI restriction sites, either with or without the first 33 nucleotides of its 3′UTR, upstream and in-frame with firefly luciferase (FLuc) in the pcDNA3.1/myc-His B vector. A linker sequence (5′-GGC GGC TCC GGC GGC TCC CTC GTG CTC GGG-3′) was inserted between *RPL36A* and FLuc. The stop codon of the dORF and the start codon of FLuc were excluded to allow continuous translation into FLuc. For GFP-based constructs, FLuc was replaced with GFP lacking a start codon and fused to two tandem HA tags (2×HA). To examine the effect of stress conditions on dORF translation, a construct containing the canonical *RPL36A* 5′UTR and CDS followed by iUTR, dORF, and FLuc was generated.

The construct used for the IRES assay contained *Renilla* luciferase (RLuc) at the 5′ end and firefly luciferase (FLuc) at the 3′ end. As a positive control, a 572-nucleotide encephalomyocarditis virus (EMCV) IRES sequence was cloned between the HindIII and EcoRI sites. As a negative control, a 267-nucleotide non-specific sequence derived from the coding region of La, lacking known IRES activity, was cloned between the same restriction sites. Both control plasmids were kindly provided by Prof. Saumitra Das’s laboratory at the Indian Institute of Science (50). For the test construct, the distal 75 nucleotides of the *RPL36A* coding sequence, followed by the canonical stop codon and the 15-nucleotide intercistronic region (iUTR), were cloned between RLuc and FLuc using the BamHI and XhoI restriction sites.

To investigate the role of leaky scanning, a stable hairpin sequence (5′-GCG GTC CAC CAC GGC CGA TAT CAC GGC CGT GGT GGA CCG CAA-3′) was used as a cis-acting molecular roadblock (54). The hairpin sequence was inserted either immediately upstream (HindIII site) or downstream (BamHI site) of the *RPL36A* coding sequence (CDS). As a control, a length-matched sequence lacking predicted secondary structure (5′-CAA CAA CAA CAA CAA CAA CAA CAA CAA CAA CAA CAA CAA CAA-3′) was used in place of the hairpin sequence, as CAA repeats are not expected to form stable RNA secondary structures under these conditions (54).

To investigate the effect of the dORF on the canonical *RPL36A* ORF, the full-length *RPL36A* coding sequence (CDS) together with its 3′UTR was cloned downstream and in-frame with an N-terminal FLAG-HA tag in the pcDNA3.1/myc-His B vector using the NotI and PciI restriction sites. The endogenous start codon of *RPL36A* was removed, and an AUG start codon was introduced upstream and in-frame with the FLAG-HA tag to drive translation of the tagged protein.

All mutations were generated using a PCR-based site-directed mutagenesis method. The same strategies were used to construct plasmids related to *RPS3* studies.

### Cell culture

HEK293 and HeLa cells were cultured in Dulbecco’s Modified Eagle Medium (DMEM; HiMedia, AL007A) supplemented with 10% fetal bovine serum (FBS; Gibco, A5256701) and 1% antibiotic solution containing penicillin (10,000 units/ml) and streptomycin (10,000 μg/ml) (Sigma, P4333). Cells were maintained at 37°C in a humidified incubator with 5% CO₂. Opti-MEM (Gibco, 11058021), glucose-free DMEM (Gibco, 11966025) supplemented with 1 mM sodium pyruvate (Sigma, S8636), and amino acid-free DMEM (US Biological, D9800-27) were used where indicated. To exclude mycoplasma contamination, cells were routinely tested using a PCR-based mycoplasma detection assay at six-month intervals (66). Cell line identity was verified by short tandem repeat (STR) profiling.

### Antibodies and reagents

The following primary antibodies were used in this study: anti-L36a (1:1,500; Santa Cruz Biotechnology, sc-100831), anti-HA (1:2,000; Sigma, 11867423001), anti-puromycin (1:500; Developmental Studies Hybridoma Bank, PMY-2A4), and anti-actin (1:30,000; Sigma, A3854). Horseradish peroxidase (HRP)-conjugated secondary antibodies were obtained from Jackson ImmunoResearch Laboratories (115-035-003 and 712-035-153). Puromycin (P8833) and MTT [3-(4,5-Dimethylthiazol-2-yl)-2,5-Diphenyltetrazolium bromide; M5655] were purchased from Sigma. Lipofectamine 2000 was obtained from Invitrogen. hsa-miR-5701 mimic (HY-R01783), microRNA mimic negative control (HY-R04602), hsa-miR-5701 inhibitor (HY-RI01783), and microRNA inhibitor negative control (HY-RI04602) were purchased from MedChemExpress.

### Transfection

Cells were seeded in 24- or 6-well plates and cultured overnight to reach ∼70–80% confluency. Transfections were performed using Lipofectamine 2000 (Invitrogen) according to the manufacturer’s instructions, with 500 ng plasmid DNA per well in 24-well plates or 2,000 ng per well in 6-well plates. For luminescence-based assays using FLuc reporter constructs, a plasmid expressing *Renilla* luciferase (RLuc; 50 ng per well in a 24-well plate) was co-transfected to normalize for transfection efficiency. Cells were harvested 24 h after transfection for downstream analyses. To investigate the effect of miR-5701, cells were transfected with miR-5701 mimic, miR-5701 inhibitor, or corresponding negative control mimic/inhibitor oligonucleotides (MedChemExpress) at a final concentration of 100 nM per well in a 24-well plate using Lipofectamine 2000. Cells were harvested after 24 h for RNA isolation and RT-PCR analysis, and after 96 h (inhibitor-treated cells) or 120 h (mimic-treated cells) for western blot analysis.

### Luminescence-based assays

Reporter-transfected cells were lysed using 1× Passive Lysis Buffer (Promega), and firefly luciferase (FLuc) and *Renilla* luciferase (RLuc) activities were measured using the Dual-Luciferase Reporter Assay System (Promega) on a GloMax Explorer luminometer (Promega). Relative luciferase activity was calculated as the ratio of FLuc activity to RLuc activity.

### Translation of the dORF under stress conditions

HEK293 cells were co-transfected with the 5′UTR-containing reporter construct and the RLuc control plasmid. Four hours after transfection, the culture medium was replaced with either complete DMEM, serum-free DMEM, glucose-free DMEM, or amino acid-free DMEM. Cells were incubated for an additional 20 h, after which relative luciferase activity was measured as described above.

### *In vitro* transcription and translation

*In vitro* transcription reactions were performed using 1 μg of AgeI-linearized plasmid. Following transcription, samples were treated with RNase-free DNase I (Thermo Scientific) at 37°C for 30 min to remove residual template DNA. The *in vitro*-transcribed RNA was purified by lithium chloride precipitation (Sigma, L9650). RNA concentration was determined using a NanoDrop 2000 spectrophotometer (Thermo Scientific). RNA integrity was assessed by electrophoresis on a native agarose gel using the RiboRuler High Range RNA Ladder (Thermo Scientific, SM1823) after incubating 200 ng RNA with a formamide-containing RNA loading dye (Thermo Scientific, R0641). *In vitro* translation was performed overnight at 30°C using 1.5 μg of the *in vitro*-transcribed RNA and Rabbit Reticulocyte Lysate (Promega L4960) according to the manufacturer’s instructions. Luciferase activity was subsequently measured as described above.

### SDS-PAGE and western blotting

After harvesting, cells were washed with ice-cold 1× phosphate-buffered saline (PBS) and lysed in radioimmunoprecipitation assay (RIPA) buffer containing 50 mM Tris-HCl (pH 8.0), 150 mM NaCl, 1 mM sodium deoxycholate, 0.1% SDS, and 1% Nonidet P-40, supplemented with 1× protease inhibitor cocktail (Promega, G6521). Total protein concentration was determined using Protein Assay Dye Reagent (Bio-Rad, 5000006) according to the manufacturer’s instructions. Cell lysates were mixed with 5× sample buffer [250 mM Tris-HCl, 10% SDS, 30% (v/v) glycerol, 0.05% (w/v) bromophenol blue, and freshly added 5% (v/v) β-mercaptoethanol] and boiled at 95°C for 5 min. Proteins were resolved by SDS–PAGE using 15% polyacrylamide gels and transferred onto 0.2 or 0.45 μm polyvinylidene difluoride (PVDF) membranes (Immobilon-P, Merck Millipore) using a Mini Trans-Blot Cell (Bio-Rad). Membranes were blocked with 5% skimmed milk for 1 h at room temperature and washed three times with PBS containing Tween-20 (PBST). Blots were incubated with the indicated primary antibodies overnight at 4°C, followed by three washes with PBST and incubation with horseradish peroxidase (HRP)-conjugated secondary antibodies for 2 h at room temperature. After three additional washes with PBST, signals were detected using Clarity Western ECL Substrate (Bio-Rad, 170-5061) or SuperSignal West Femto Maximum Sensitivity Substrate (34095) in a Chemidoc (Bio-Rad). Band intensities were quantified using ImageJ. Background-subtracted intensities of proteins of interest were normalized to the loading control (actin), and identical-sized regions were used for densitometric analysis across all samples.

### RNA isolation and quantitative real-time PCR (qRT-PCR)

Total RNA was isolated using RNAiso Plus (TaKaRa, 9108) according to the manufacturer’s instructions. RNA concentration and purity were determined using a NanoDrop 2000 spectrophotometer (Thermo Scientific), and RNA integrity was assessed by agarose gel electrophoresis. First-strand cDNA synthesis was performed using oligo(dT) primers (Bioserve Biotechnologies) and RevertAid Reverse Transcriptase (Thermo Scientific, EP0441). qRT-PCR was carried out using TB Green Premix Ex Taq II (TaKaRa, RR820) in 96-well Hard-Shell PCR Plates (Bio-Rad) on a Bio-Rad CFX Opus 96 Real-Time PCR system. The cycling conditions were: 95°C for 30 s, followed by 35 cycles of 95°C for 5 s, 55°C for 30 s, and 72°C for 30 s, with a final extension step at 72°C for 5 min. Melting curve analysis was performed after each run. Primer specificity was verified by agarose gel electrophoresis of the amplified products. Relative gene expression levels were calculated using the 2^−ΔΔCt method.

For miRNA expression analysis, total RNA or small RNA-enriched fractions were isolated using the miRNeasy kit (Qiagen, 217084) according to the manufacturer’s protocol. First-strand cDNA synthesis for miR-5701 was performed using the RT11-miR-5701 primer (Bioserve Biotechnologies), whereas cDNA synthesis for the normalization control U6 was carried out using random hexamers (Invitrogen). miRNA qRT-PCR was performed using PowerUp SYBR Green Master Mix (Applied Biosystems, A25742) on the Bio-Rad CFX Opus 96 system (67). The cycling conditions were: 50°C for 2 min, 95°C for 2 min, followed by 40 cycles of 95°C for 15 s, 53.1°C for 15 s, and 72°C for 1 min. Melting curve analysis and agarose gel electrophoresis were used to confirm amplification specificity.

Primer sequences (5′→3′):

Exogenous *RPL36A*:

CCTTATGACGTGCCCGATTAC and GTCATAACGCCGCTTTCCCTGG

Endogenous *RPL36A*:

GATAGCGCTCACGCAAGCATG and GTCATAACGCCGCTTTCCCTGG

*FGFR2*:

TGACATTAACCGTGTTCCTGAG and TGGCGAGTCCAAAGTCTGCTAT

*FLuc*:

CAACTGCATAAGGCTATGAAGAGA and ATTTGTATTCAGCCCATATCGTTT

*Actin*:

CACCAACTGGGACGACAT and ACAGCCTGGATAGCAACG

miRNA common:

TGTCAGGCAACCGTATTCACC and CGTCAGATGTCCGAGTAGAGG

RT11-miR-5701:

TGTCAGGCAACCGTATTCACCGTGAGTGGTAATCAGAACGT

Short miR-5701 REV:

CGTCAGATGTCCGAGTAGAGGGGGAACGGCGTTATTGTCACGTT

*U6*:

CTCGCTTCGGCAGCACAT and TTTGCGTGTCATCCTTGCG

### CRISPR-Cas-mediated genomic deletion of the proximal 3′UTR of *RPL36A*

The proximal 3′UTR region of *RPL36A* was targeted using two sgRNAs: 5′-AGTGTCATCTTTTATTATGA-3′ and 5′-ATGAAGACAATAAAATCTTG-3′ (Bioserve Biotechnologies). sgRNAs were designed using Benchling and cloned into the pX330-SpCas9-NG plasmid. A turboGFP cassette was inserted into the KpnI site of the plasmid to enable fluorescence-activated cell sorting (FACS)-based enrichment of transfected cells. HeLa cells at ∼70-80% confluency in 6-well plates were co-transfected with the two sgRNA-expressing plasmids (1,000 ng each) using Lipofectamine 2000 (Invitrogen). Forty-eight hours after transfection, the top 5% GFP-positive cells were sorted using a BD FACSAria Fusion Flow Cytometer (BD Biosciences) and seeded at a density of one cell per well in 96-well plates. Expanded single-cell clones were screened for genomic deletion by PCR amplification of the target region using primers flanking the edited locus (5′-GGTGAGTATTTTAGGAACAG-3′ and 5′-GAATTGTATTGATGGCTCTAGC-3′; Bioserve Biotechnologies). PCR products were validated by Sanger sequencing to confirm the deletion. The resulting *RPL36A* dORF deletion clone of HeLa cells was heterozygous for the deletion.

### Ribopuromycylation assay

Wild-type and *RPL36A* ΔdORF HeLa cells were seeded at equal densities (1×10^6 cells per well in 6-well plates). 7 h after seeding, cells were washed once with 1× PBS and cultured in serum-free DMEM for 12 h. Cells were then incubated with puromycin (91 μM) for 10 min at 37°C by replacing the medium with fresh serum-free medium containing puromycin (68). Following puromycin treatment, cells were harvested, lysed, and western blotting was performed to detect puromycylated proteins using anti-puromycin antibody.

### Prediction of microRNA-binding sites and miRNA expression analysis

Predicted microRNA-binding sites within the *RPL36A* mRNA were identified using miRDB, an online database for miRNA target prediction and functional annotation (56). The first 100 nucleotides of the *RPL36A* 3′UTR, encompassing the dORF region, were used as the input sequence for analysis. Expression profiles of hsa-miR-6740-3p, hsa-miR-511-5p, and hsa-miR-5701 in HEK293 cells were analyzed using the web-based DIANA-miTED database (57).

### Statistics

Paired or unpaired Student’s t-test was used to calculate statistical significance, as described in the figure legends. Welch’s correction was applied to samples that demonstrated differences in variance.

## ACKNOWLEDGEMENTS

This work was supported by funding from the Swarnajayanti Fellowship (DST/SJF/LSA-04/2019-20) awarded by the Department of Science and Technology (DST). Additional support was provided by the STARS grant from the Ministry of Education (STARS/APR2019/BS/328/FS), DBT’s Genome Engineering Technology Program (BT/PR38405/GET/119/309/2020), the Indian Council of Medical Research, Anusandhan National Research Foundation, the Blockchain Foundation of India, the EMBO Global Investigator Network, DST Funds for Improvement of Science and Technology infrastructure, and the Institute of Eminence funds allocated by the Ministry of Education to the Indian Institute of Science. JG acknowledges a doctoral fellowship from the Council of Scientific and Industrial Research.

