## Supplemental Figures for "Leaky scanning translates a conserved ORF in the 3′UTR of *RPL36A* and regulates the expression of ribosomal protein L36a"

### Figure S1

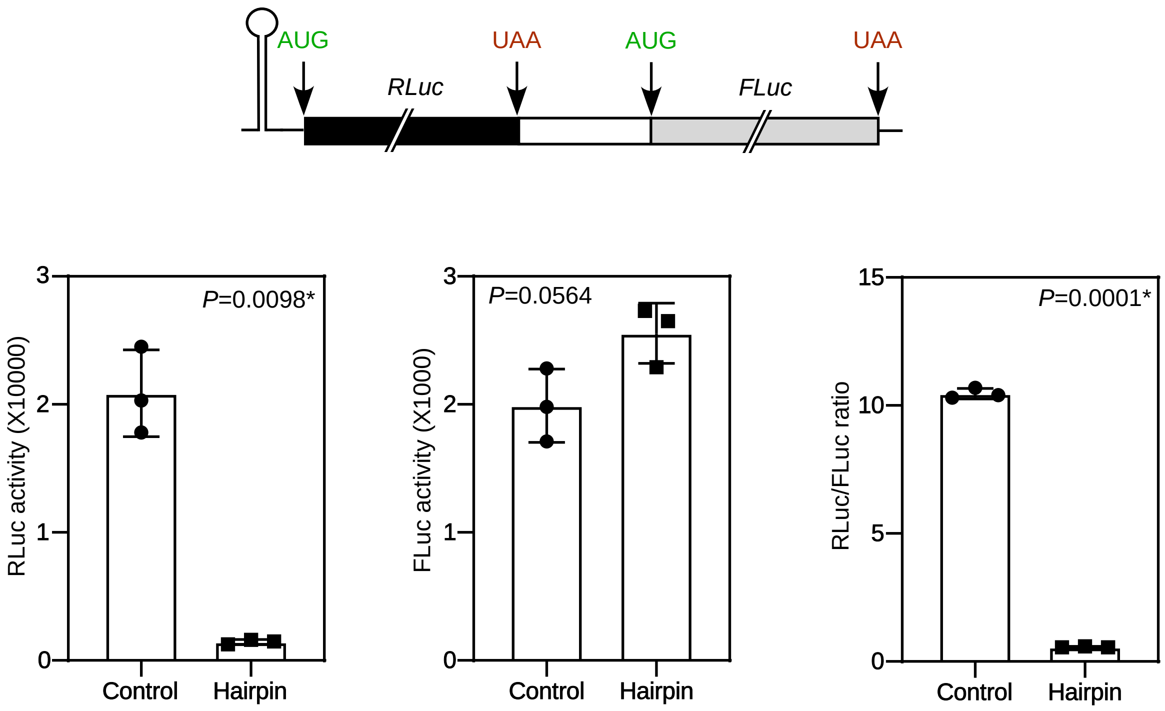


**Figure S1. Inhibition of cap-mediated translation by a stem-loop forming sequence.**
The schematic shows the dual-luciferase construct used to demonstrate the effect of the hairpin structure on cap-mediated translation. Expression of *Renilla* luciferase (RLuc) was mediated by the 5' cap, and the firefly luciferase (FLuc) was mediated by an IRES. A 42-nucleotide hairpin-forming sequence or a length-matched control sequence was inserted upstream of RLuc. The plasmid was transfected into HEK293 cells, and the activity of luciferase was measured as explained in Methods. The graphs show RLuc activity, FLuc activity, and their ratios. Bars in all graphs represent mean ± SD, n = 3. Statistical significance was determined using a two-tailed unpaired Student's t-test (* Welch's correction was applied).

### Figure S2

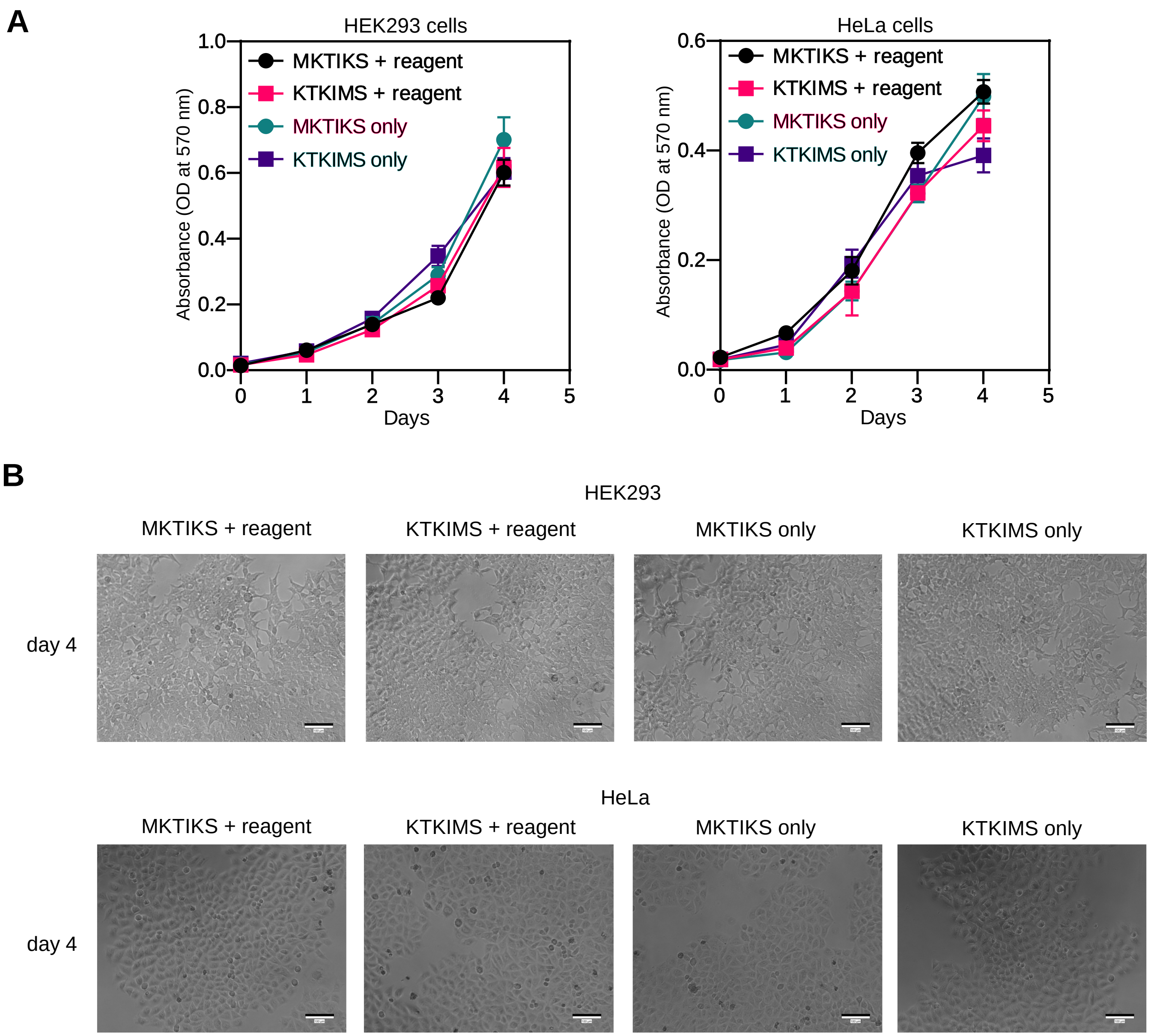


**Figure S2. The peptide encoded by the *RPL36A* dORF does not exert a detectable effect on cells.**
(A) Proliferation of cells treated with the indicated peptides.
(B) Images of the peptide-treated cells after 4 days of treatment. Images were acquired using an IXplore IX73 inverted microscope (Olympus) with a 10× objective lens.
The peptide encoded by the *RPL36A* dORF (MKTIKS) and a scrambled control peptide (KTKIMS) were synthesized by Pentavalent Bio Sciences Pvt. Ltd., India. Peptides were reconstituted in sterile water prior to use. HEK293 and HeLa cells at ~70–80% confluency in 24-well plates were treated with peptides (500 nM) using Xfect Peptide Transfection Reagent (TaKaRa) according to the manufacturer's instructions or without any transfection reagent. Four hours after transfection, the medium was replaced with complete growth medium, and cells were incubated for an additional 2 h. For cell proliferation assays, peptide-transfected cells were trypsinized and counted, and equal numbers of cells (~2,000 cells per well) were seeded into 96-well plates. Cell proliferation was monitored over 4 days using the MTT assay. At the indicated time points, 50 µg MTT [3-(4,5-Dimethylthiazol-2-yl)-2,5-diphenyltetrazolium bromide] was added to each well, followed by incubation at 37°C for 2 h. The medium was then removed, and 100 µl DMSO was added to each well to dissolve the formazan crystals. After incubation for 15 min, the dissolved reaction mixture was transferred to a fresh 96-well plate, and absorbance was measured at 575 nm using a VERSAmax microplate reader (Molecular Devices).

### Figure S3

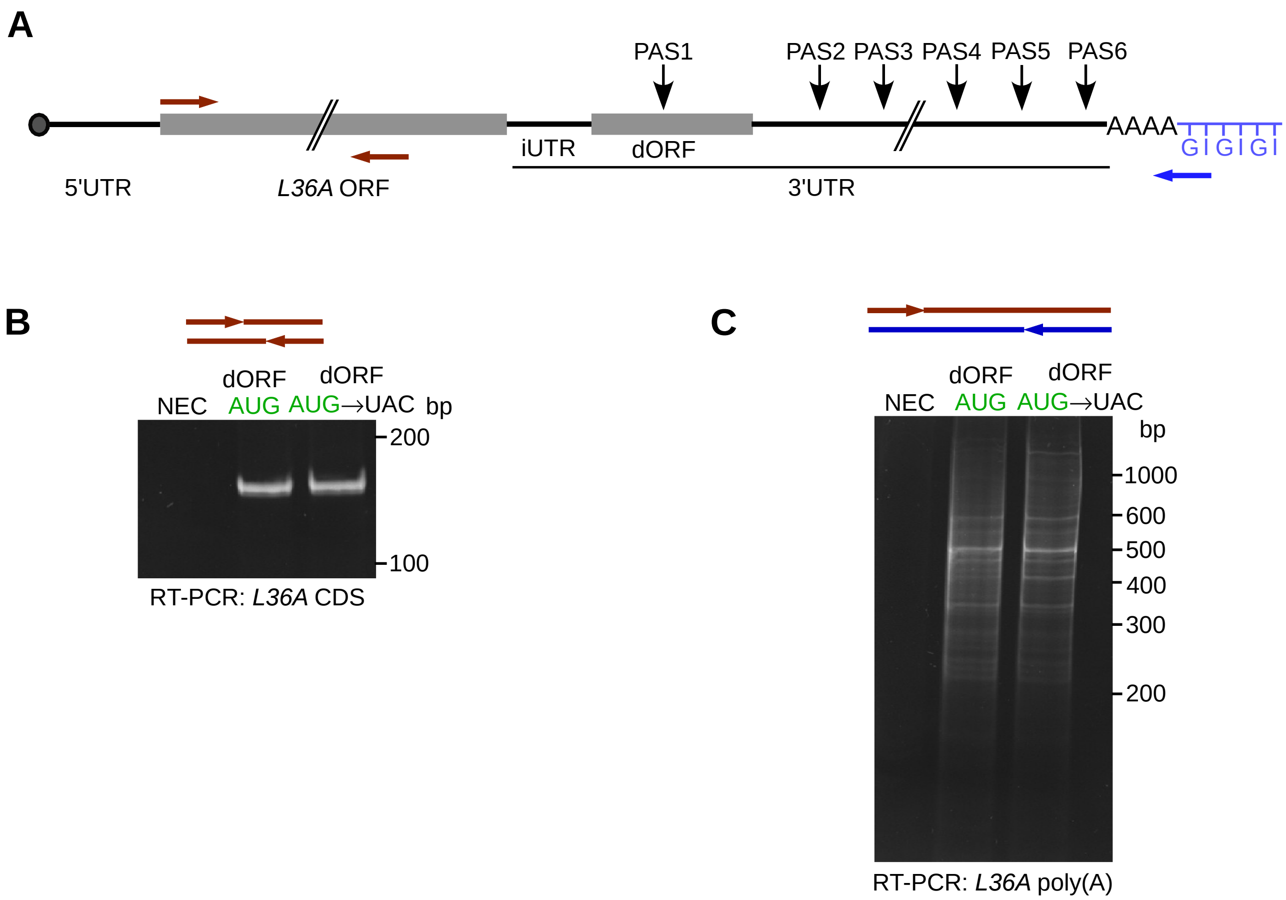


**Figure S3. Disruption of *RPL36A* dORF translation does not alter the choice of polyadenylation site.**
(A) Schematic of the exogenous *RPL36A* transcript showing the canonical ORF, the dORF, and the predicted alternative polyadenylation sites (PAS) within the 3'UTR. Positions of the primers used for RT-PCR and G/I-tailing-based poly(A) site analysis are indicated.
(B) RT-PCR analysis of exogenous *RPL36A* transcripts from cells transfected with constructs carrying *RPL36A* with either the wild-type or start-codon-mutated dORF, using the indicated primer pairs.
(C) G/I-tailing assay used to assess polyadenylation-site usage of exogenous *RPL36A* transcripts isolated from cells transfected with constructs carrying *RPL36A* with either the wild-type or mutant dORF. A 5'-phosphorylated DNA adaptor oligonucleotide (5'-GIGIGIGIGIGIGIGIGIGIGIGIG-3') was ligated to the 3' hydroxyl termini of total RNA using T4 RNA ligase (Thermo Scientific, EL0021). Following adaptor ligation, first-strand cDNA synthesis was performed using the adaptor- and poly(A)-complementary primer C10T7 REV (5'-CCCCCCCCCCTTTTTTT-3') with RevertAid Reverse Transcriptase. Subsequently, two RT-PCR reactions were performed for each sample. One reaction amplified only the *RPL36A* coding sequence (CDS) using a CDS-specific primer pair (shown in B). The second reaction (shown in C) amplified full-length RPL36A transcripts using a CDS-specific forward primer and the adaptor/poly(A)-complementary C10T7 REV primer with Phusion High-Fidelity DNA Polymerase. PCR cycling conditions were: 98°C for 3 min, followed by 20 cycles of 98°C for 30 s, 57°C for 50 s, and 68°C for 2 min, with a final extension at 68°C for 10 min. PCR products were resolved on 8% Tris-Borate-EDTA (TBE) gels, stained with ethidium bromide, and visualized using a Bio-Rad ChemiDoc imaging system.
Data shown are representative of three independent experiments.
